# Early Tactile Stimulation Influences the Development of Alzheimer Disease in Gestationally Stressed APP ^NL-G-F^ Adult Offspring ^*NL-G-F/NL-G-F*^ mice

**DOI:** 10.1101/2022.02.28.482233

**Authors:** Shakhawat R. Hossain, Hadil Karem, Zahra Jafari, Bryan E. Kolb, Majid H. Mohajerani

**Author notes:** Majid H. Mohajerani and Bryan E. Kolb are corresponding authors, contributed equally to the manuscript, and share the senior author position for this paper.

## Abstract

Alzheimer Disease (AD) is associated with cerebral plaques and tangles, reduced synapse number, and shrinkage in several brain areas and these morphological effects are associated with the onset of compromised cognitive, motor, and anxiety-like behaviours. The focus of this study was to examine the effect of neonatal tactile stimulation on AD-like behavioural and neurological symptoms on APP ^*NL-G-F/NL-G-F*^ mice, a mouse model of AD. Our findings indicate that neonatal tactile stimulation improves cognition, motor skills, and anxiety-like symptoms in both gestationally stressed and non-stressed adult APP mice and that these alterations are associated with reduced Aβ plaque formation. Thus, tactile stimulation appears to be a promising non-invasive preventative strategy for slowing the onset of dementia in aging animals.

## Introduction

Alzheimer disease (AD) is a neurodegenerative brain disorder that causes cognitive and motor skill deficits combined with a lack of motivational, social, and emotional behaviours. These behavioural symptoms are associated with the neural symptoms such as: shrinkage of the cerebral cortex, hippocampus, and basal ganglia (Pini et al., 2016); formation of extracellular Aß plaques and intracellular tau phosphorylated proteins (Marcello et al., 2015); synaptic loss (Hamos et al., 1989); and disrupted gamma oscillations (Iaccarino et al., 2016) in the brain. There are many identified risk factors for AD such as depression, auditory stress as a form of noise pollution, diabetes, anoxia, high blood pressure, low education, smoking, alcohol, obesity, brain injury, and physical inactivity to name few (Alzheimer’s Society Canada, 2019).

Auditory/noise stress has been identified as strong risk factor of AD. Research on humans has demonstrated that nighttime noise levels, and distance from major roads are highly associated with AD (Carey et al., 2018). Research on mice has demonstrated that exposure to noise stress during the prenatal (Jafari et al., 2018; 2019) or postnatal periods (Jafari et al., 2017; Cheng et al., 2011) has a detrimental effect on brain structures, neural networks, and related cognitive behavior in offspring and adult mice respectively (Jafari et al., 2019; Arnsten, 2015). Gestational auditory stress has proven to be harmful for infant brain and behavioural development and aggravates cognitive deficits and Aβ plaque formation in APP mice, a mouse model of AD (Jafari et al., 2019). In addition, prenatal stress has proven to alter the organization of neural circuits in the neocortex and hippocampus in rats (Mychasiuk et al., 2012). There is plenty of evidence from clinical research on humans showing that people living in major cities (Astell-Burt & Feng, 2018) and closer proximity to heavy traffic noise (Chen et al., 2019) have higher chance of being diagnosed with AD/dementia along with other cognitive impairment disorders (Tzivian et al., 2016). Previous research in our lab has demonstrated the adverse effects of auditory stress on the brain and behaviour in second generation of mouse model of AD (Jafari et al., 2017; Mehla et al., 2019; Karem et al., 2021). In this study, we aimed to explore the role of early tactile stimulation (TS) in mitigating the adverse effects of gestational auditory stress, as well as AD-like symptoms in adult APP offspring.

TS has beneficial impact on brain and behavioral development. Research using rodents has shown that TS aided recovery from brain injury (Gibb et al., 2010). One mechanism may be that TS releases fibroblast growth factor-2 (FGF-2) from the skin, which stimulates neurogenesis, cellular proliferation, survival, migration, and neuronal differentiation. Research on premature human infants showed that TS improves the development of their cognitive and motor skills (Field et al., 1986) and may stimulate neuroplasticity later in life. Laboratory research on rodents has also established that the improved cognitive and motor skills are associated with: increased FGF-2 (Comeau et al., 2007), BDNF (Antoniazzi et, al., 2016), acetylcholine (Dudar et al., 1979), and synaptic plasticity (Kolb & Gibb, 2010; 2011) in the brain.

The effect of TS is strongest when it is applied during the early infantile period. Therefore, we designed this research project in which we applied TS from postnatal day 3-18 and observed the behavioural improvements at two months in one group of APP adult offspring and in another group at six months. Our predictions were: 1) early TS would mitigate the adverse effects of gestational auditory stress in adult APP offspring; 2) early TS would improve the cognitive, motor, and anxiety-like behaviours in adult APP offspring; and, 3) early TS would reduce the development of Aß plaques in both APP and APP-GS adult offspring.

## Methods and Materials

### Animals

APP^NL-G-F/NL-G-F^ (amyloid ß-protein precursor), Alzheimer disease transgenic mice carrying Swedish (NL), Arctic (G), and Beyreuther/Iberian (F) mutations were used in this research project. Twenty-four 8 week old female APP^NL-G-F/NL-G-F^ mice were individually mated with twenty-four male APP^NL-G-F/NL-G-F^ mice in standard shoe-box cages at 4:00 pm. For the recording of gestational length, a former protocol was followed (Jafari, et al. 2017). Upon the confirmation of ++ genotyping of the offspring using the tail snipping method, sixty female APP^NL-G-F/NL-G-F^ offspring and sixty male APP^NL-G-F/NL-G-F^ offspring were used in this project (see Figure 1). Mice were housed in Canadian Center for Behavioral Neuroscience (CCBN) vivarium, and all the behavioral, brain anatomical and physiological tests and analyses were approved by the Animal Welfare Committee at the University of Lethbridge. The mice were maintained on a 12-hour light and 12 hours dark cycle in a 21°C temperature controlled room in the vivarium and training and behavioral testing was performed by the same experimenter during the light phase.

**Figure 1.**
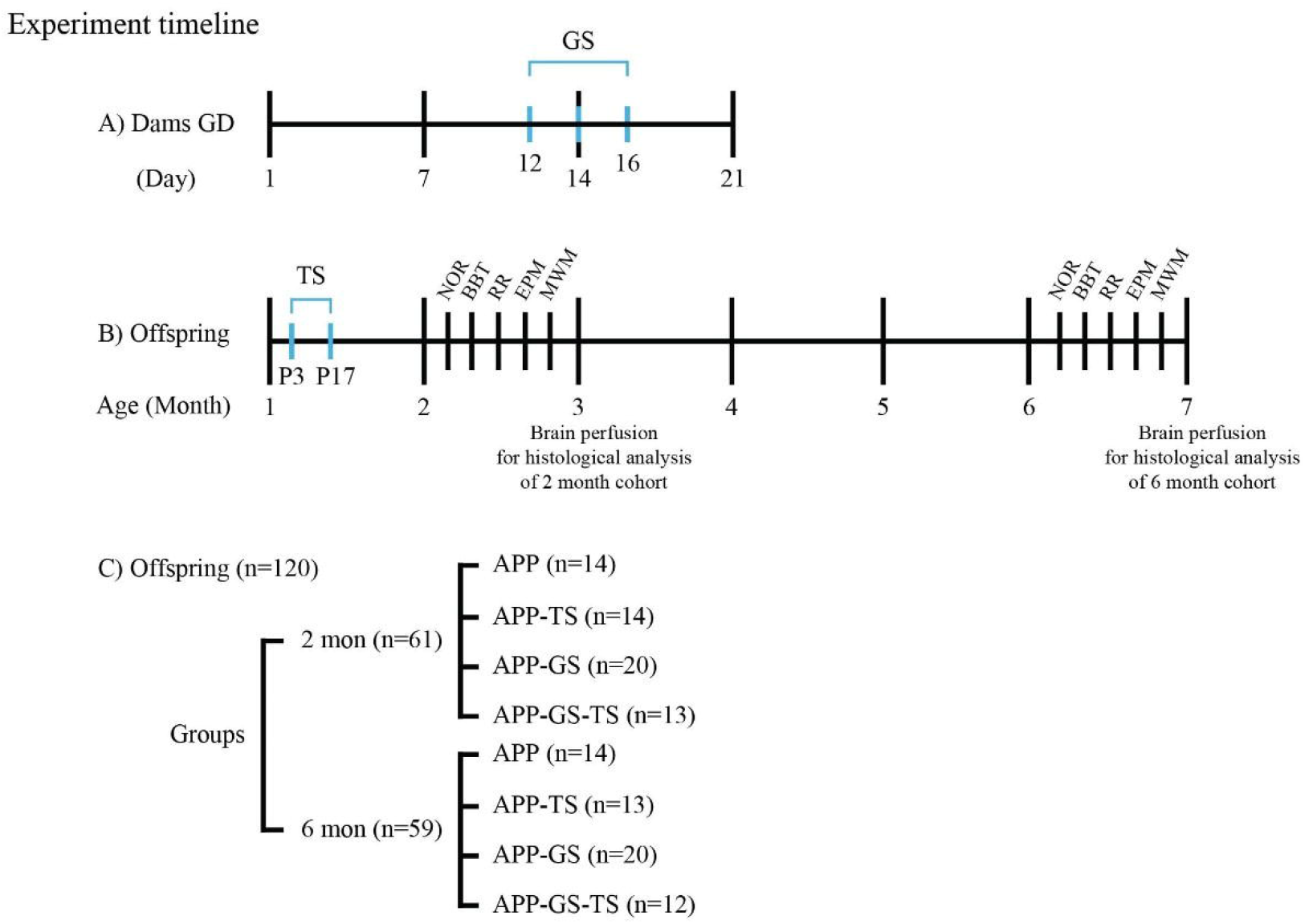
(A) Timeline for gestational period and half of the pregnant mice received GS on GD 12, 14, and 16 for 24 hours per day. (B) The timeline for offspring shows half of the pups received TS during Postnatal day 3-17, behavioural measurement for 2- and 6-months cohorts respectively, and brain perfusion at 3 and 7 months for 2- and 6-months cohorts respectively. (C) The experimental groups and total number of mice in each group consisting both male and female mice.

### Stress Paradigm

Half of the pregnant mice were exposed to acute auditory sound (AS) stress for 24 hours every other day with an intensity of 90 dB during gestational day (GD) 12-16. The AS, consisted of an intermittent 3000 Hz frequency, was programmed by using MATLAB software and played the same repetitive sound for 1 second for 5400 times with an interval of 15 seconds between 2 sounds (Haque et al. 2004; Jafari, Faraji, et al. 2017). The reasons for using 3000 Hz intermittent frequency are that 3000 Hz is similar to traffic and environmental noise (Chang et al. 2014) and is easily audible to mice (Heffner and Heffner, 2007).

### Experimental Design

The offspring from both gestational stress (GS) and no gestational stress (NGS) groups of APP^NL-G-F/NL-G-F^ mice were randomly assigned to two cohorts: 2 months and 6 months. Offspring from each of 2- and 6-months cohorts were further assigned to four different groups consisting of APP, APP-Tactile Stimulation (APP-TS), APP-Gestational Stress (APP-GS), and APP-Gestational Stress-Tactile Stimulation (APP-GS-TS), as per the following table. Each group consists of a minimum 4 male and 4 female mice using a cross litter design to avoid possible litter effects.

### Tactile Stimulation Procedure

Half of the offspring from both GS and NGS mice groups received TS for 15 minutes by gently brushing the body of the offspring using a soft sweeper broom at postnatal day (PD) 3-17, for 15 minutes with a frequency of 3 times a day for 15 days (8 am, 12 pm, and 4 pm). The offspring were weaned at PD 21 and housed with a group of 3-4 mice per cage until 3 and 7 months in the same animal room per 2- and 6- months cohorts respectively.

### Behavioral Tests

All the mice in both 2- and 6-months cohorts performed several behavioral tests at 2 and 6 months, respectively, to measure the effect of TS on cognitive and motor functions and anxiety-like behaviours. Balance beam (BB), rota-rod (RR), novel object recognition (NOR), activity box (AB), elevated plus maze (EPM), and the Morris water task (MWT) were conducted by the same examiner with an alternating order of animals (see Figure 1). The mice in 2- and 6-months cohorts were perfused at 3 and 7 months respectively, at the completion of the behavioral testing.

### Novel Object Recognition (NOR) Test

The NOR test was used to measure the short-term memory in the mice. Each mouse was placed in the same open field arena of 47cm x 50cm x 30cm with 2 similar objects for 5 minutes. Following a 3-minute break, each mouse was exposed to one old and a novel object and the activity was filmed for 3 minutes. The time (seconds) spent with each old and novel object was manually scored for analysis (Jafari et al., 2017). The discrimination index (DI) was calculated by subtracting time spent with old object from the time spent with the novel object divided by the total time spent with both novel and old objects (Ennaceur & Delacour, 1988).

### Morris Water Task (MWT)

The MWT task was performed to measure the spatial navigation abilities in the mice. Each mouse was placed in a 153cm diameter pool, filled with water (23-25°C), which was located in a room with distal cues. The pool was virtually subdivided into 4 quadrants with starting points at north, west, east, and south and a ~1.0 cm submerged platform was placed in one fixed quadrant during all 8 training days. Non-toxic white tempura paint was added to the pool water to make the water opaque to make the platform invisible. Each mouse was trained with 1 trial per day from each quadrant for 8 consecutive days (Water2100 Software vs.7, 2008). During each trial, the mouse was placed in the pool and each trial was stopped either once the mouse reached the platform, or if the mouse was unable to find the platform in 60 seconds. Data were recorded using an automated tracking system (HVS Image, Hampton, U.K.) and swim time (sec), swim speed (m/s), and swim distance (m) were calculated for analysis. On day 9, a probe trial was conducted, during which the platform was removed, and each mouse was allowed to swim freely for 60 seconds. For the analysis of the probe trial, the time spent in the quadrant where the platform was located during training days was measured (Jafari et al., 2017).

### Balance Beam (BB) Test

The BB test was performed to measure the motor skills of mice. Each mouse was trained to traverse across a 1 cm diameter, 100 cm long beam, which was 50 cm above a foam pad to cushion falling mice, to reach an escape box. Each mouse was trained to traverse the beam for 3 successful trials on day 1. Traverse activity of each mouse was recorded for 3 trials on day 2 and manually scored for the mean latency (sec), distance travelled (cm), and number of foot slips for analysis (Jafari et al., 2018; Tamura et al., 2012).

### Rotarod (RR) Test

The RR test measures the motor skills and the strength of gait in each mouse. On day 1, mice were trained to walk on an automated four lanes RR treadmill (ENV-575 M Mouse, Med Association Inc). On day 2, each mouse was placed on the RR treadmill at 8rpm and 16rpm constant speed and at a 4-40 rpm alternating speed. The time (sec) each mouse was able to stay on the RR treadmill was recorded for 3 trials on day 2 for analysis (Brooks and Dunnett, 2009).

### Elevated Plus Maze (EPM) Test

The EPM measures anxiety-like behaviour in mice. The EPM apparatus was constructed from black Plexi-glass and had two closed arms of 10 cm wide and 4 cm long and two open arms of 5 cm wide and 27 cm long. Each mouse was placed in the center of the EPM facing a closed arm and filmed for 5 minutes. The time spent in the open arms (seconds) and in closed arms (seconds), and number of entries to open arms and to closed arms were scored manually for each mouse (Jafari et al., 2017). The EPM ratio was calculated by subtracting the number of entries to open arms from the number of entries to closed arms divided by the total number of entries to both open and closed arms (Jafari et al., 2018).

## Results

In both 2- and 6-months cohorts there were four different groups: APP, APP-TS, APP-GS, and APP-GS-TS. In this experiment 115 animals were used consistently in each behavioral test. None of the behavioral tests showed sex differences (P>05) so sex was removed as a variable in the statistical analyses. Two-way ANOVA was done for 2 months and 6 months cohorts separately. The Bonferroni post-hoc test was used for each behavioral test, due to similar variance in each groups. Asterisks indicate *P≤0.05 or **P≤ 0.01 or ***P≤ 0.001 and partial eta squared (η2) indicates the effect size.

### Novel Object Recognition (NOR) Test

The APP-GS mice spent significantly less time with the novel object compared to all of the other experimental groups (see Figure 2). TS significantly increased the novel object exploration in both APP and APP-GS mice. The overall significant effects among all of the four groups of 2 months cohort were: novel object time (F (3, 57) = 10.623, P≤ .0001, □2= .376, power = .998) and discrimination index ratio (F (3, 57) = 13.718, P≤ .0001, □2= .437, power = 1.000). The overall significant effects among all of the four groups of 6 months cohort were: novel object time (F (3, 57) = 63.560, P≤ .0001, □2= .806, power = 1.000), old object time (F (3, 57) = 12.861, P≤ .0001, □2= .456, power = .998), and DI ratio (F (3, 57) = 14.429, P≤ .0001, □ 2= .485, power = 1.000).

**Figure 2.**
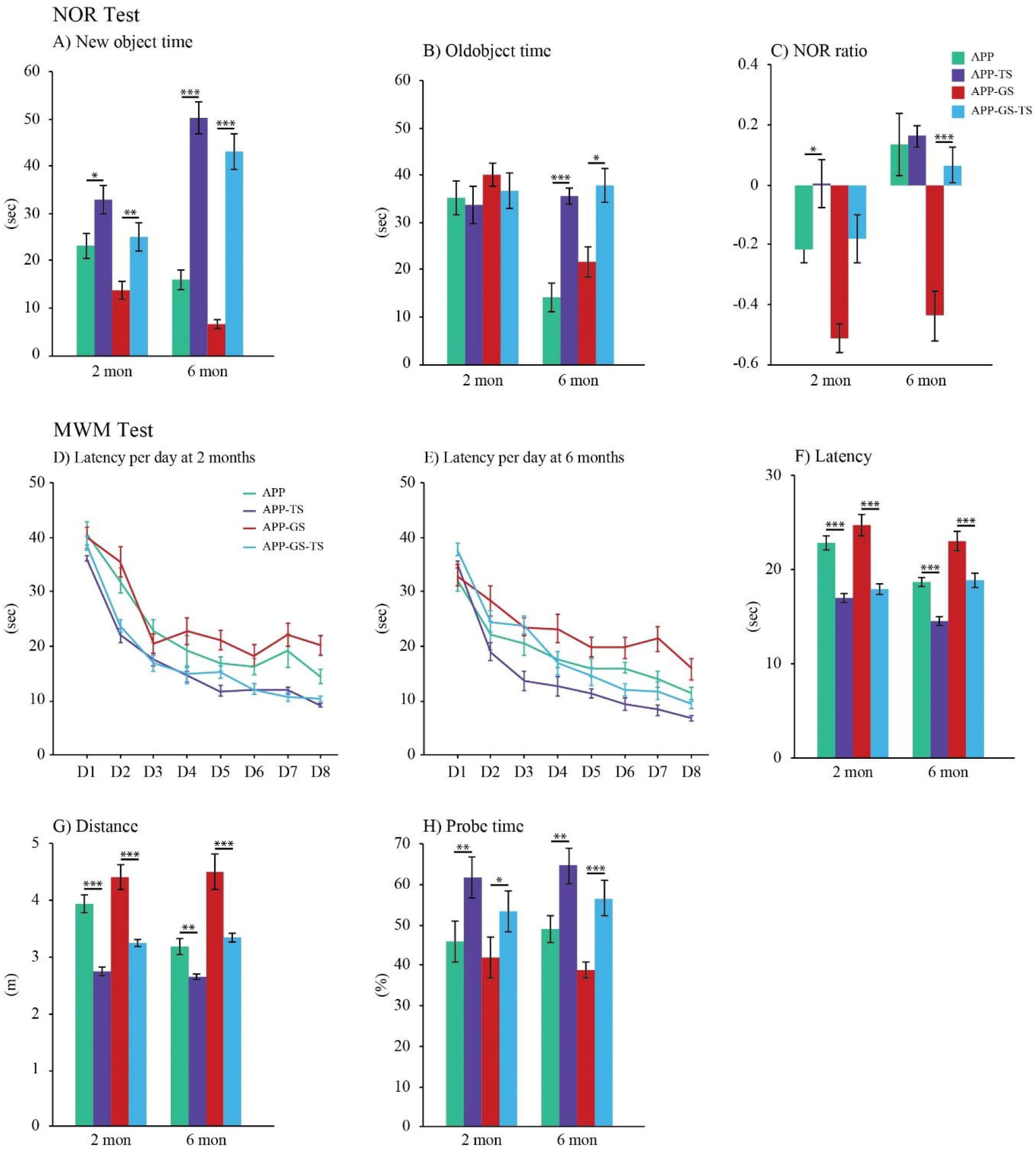
(A) The APP-TS group in both 2 and 6 months cohorts spent the highest time (sec) with the new object. (B) The time spent with the old object was not significantly different in the 2 months cohort, but the APP group spent significantly less time with old object in the 6 months cohort. (C) In both 2 and 6 months cohorts APP-GS mice showed lower preference for new object. (D & E) The progression of swim latency from Day 1-8 in both 2 and 6 months cohorts. (F) Swim latency was longer in APP-GS mice in both 2 and 6 months cohorts. (G) Swim distance was significantly higher in the APP-GS mice in both 2 and 6 months cohorts. (H) Probe time was significantly longer in APP-TS in both the 2 and 6 months cohorts.

In the 2 months cohort, the novel object time was significantly higher in APP-TS mice compared to APP (F (1, 26) = 5.972, P = .022, □2= .199, power = .650) and in APP_GS_TS mice compared to APP-GS (F (1, 26) = 12.209, P = .002, □2= .296, power = .922). Similarly, the DI ratio was significantly higher in APP-TS mice compared to APP (F (1, 26) = 5.849, P = .024, □2 = .196, power = 641) at 2 months. No significant differences were observed in the old object exploration time among all of the groups at 2 months. A Bonferroni post-hoc analysis revealed that the APP-TS group spent significantly more time with the novel object in comparison with APP-GS (P≤ .0001), and APP-GS group spent a significantly less time with novel object in comparison with APP-TS (P≤ .0001), and APP-GS-TS (P = .013) in the 2 months cohort. No significant difference was revealed from a Bonferroni post hoc analysis in the time spent with the old object among the groups. In addition, the DI ratio was highest in the APP-TS mice in comparison with APP-GS (P≤ .0001) and was lowest in the APP_GS in comparison with APP (P = .005), APP-TS (P≤ .0001), and APP-GS-TS (P = .002) as per Bonferroni post hoc analysis in the 2-months cohort.

In the 6-months cohort, the novel object time was significantly higher in APP-TS mice compared to APP (F (1, 25) = 85.778, P≤ .0001, □2 = .789, power = 1.000) and in APP-GS_TS mice compared to APP-GS (F (1, 25) = 90.527, P≤ .0001, □2 = .797, power = 1.000). In contrast, the old object time was significantly lower in APP-TS mice compared to APP (F (1, 25) = 33.046, P≤ .0001, □2 = .590, power = 1.000) and in APP-GS-TS mice compared to APP-GS (F (1, 25) = 6.532, P = .018, □2 = .221, power = .687) in the 6 months cohort. In addition, the DI ratio was significantly higher in APP-GS-TS compared to APP-GS (F (1, 25) = 25.005, P≤ .0001, □2 = .521, power = .998), but not significantly different between APP and APP-TS in the 6 months cohort. A Bonferroni post-hoc analysis revealed that the APP-TS group spent significantly more time with the novel object in comparison with APP (P≤ .0001) and the APP_GS (P≤ .0001) and APP-GS group spent significantly less time with the novel object in comparison with APP-TS (P≤ .0001) and APP-GS_TS (P≤ .0001) in 6 the months cohort. Surprisingly, APP-TS spent the highest amount of time exploring the old object in comparison with APP (P≤ .0001) and APP-GS (P = .006). The APP mice spent the lowest amount of time with the old object compared to APP-TS (P≤ .0001) and APP-GS-TS (P≤ .0001) as per Bonferroni post hoc analysis. In addition, the DI ratio was highest in the APP-TS mice in comparison with APP-GS (P≤ .0001) and was lowest in the APP-GS in comparison with APP (P≤ .0001), APP-TS (P≤ .0001), and APP-GS-TS (P≤ .0001) as per Bonferroni post hoc analysis in the 6 months cohort.

### Morris Water Task (MWT)

The APP-GS mice were significantly slower to locate the sub-merged platform, demonstrated a longer swim distance, and a reduced probe time in the target quadrant compared to all of the other groups. TS significantly improved the performances on all three measures for both of the APP and APP-GS mice (see Figure 2).

The overall significant effects among all of the four groups of 2 months cohort were: latency (F (3, 57) = 17.648, P≤ .0001, □2 = .500, power = 1.000), swim distance (F (3, 57) = 8.551, P≤ .0001, □2 = .521, power= 1.000), and probe time (F (3, 23) = 6.532, P = .001, □2= .270, power= .961). The overall significant effects among all of the four groups of 6 months cohort were: latency (F (3, 50) = 32.647, P≤ .0001, □2 = .680, power = 1.000), swim distance (F (3, 50) = 42.970, P≤ .0001, □2 = .737, power = 1.000), and probe time (F (3, 50) = 10.198, P≤ .0001, □2 = .399, power = .997).

In the 2 months cohort, the swim latency during training days was significantly decreased in the APP-TS mice compared to APP (F (1, 26) = 42.993, P≤ .0001, □2 = .642, power = 1.000) and in APP-GS-TS compared to APP-GS (F (1, 31) = 19.526, P≤ .0001, □2 = .402, power = .990). Similarly, the swim distance during the training days also was decreased significantly in the APP-TS mice relative to APP (F (1, 26) = 40.900, P≤ .0001, □2 = .630, power = 1.000) and in APP-GS-TS compared to APP-GS (F (1, 31) = 14.735, P≤ .001, □2 = .390, power = .957). In addition, during the probe day, the amount of time spent in the target quadrant was significantly higher in the APP-TS mice compared to the APP (F (1, 26) = 8.404, P =.008, □2 = .259, power = .794), in the APP-GS-TS compared to APP-GS (F (1, 31) = 6.365, P = .017, □2 = .337, power = .684). A Bonferroni post hoc analysis revealed that the APP-TS mice took significantly less time to locate the hidden platform during training days in comparison with APP (P = .001) and APP-GS (P≤ .0001) and APP-GS took significantly more time than the APP-TS (P≤ .0001) and APP-GS-TS (P≡ .0001). APP-TS mice swam the least distance compared to APP (P≤ .0001) and APP-GS (P≤ .0001) and APP-GS (P≤ .0001) swam the longest distance compared to APP-TS (p≤ .0001) and APP-GS-TS(p≤ .0001). APP-TS mice spent the highest amount of time in the target quadrant during the probe test compared to APP (P = .016) and APP-GS (P =.001) and APP-GS mice spent the least amount of time in the target quadrant compared to APP-TS (P = .001).

In the 6 months cohort, during training days the swim latency was significantly decreased in the APP-TS mice compared to APP (F (1, 25) = 39.476, P≤ .0001, □2 = .632, power = 1.000) and in APP-GS-TS compared to APP-GS (F (1, 25) = 21.243, P≤ .0001, □2 = .480, power = .993). Similarly, the swim distance during the training days was also significantly decreased in the APP-TS mice relative to APP (F (1, 25) = 14.735, P = .001, □2 = .390, power = .957) and in APP-GS-TS compared to APP-GS (F (1, 25) = 37.571, P≤ .0001, □2 = .620, power = 1.000). In addition, during the probe day, the amount of time spent in the target quadrant was significantly higher in the APP-TS mice compared to the APP (F (1, 25) = 8.222, P = .009, □2 = .263, power = .784), in the APP-GS-TS compared to APP-GS (F (1, 25) = 17.142, P≤ .0001, □2 = .427, power = .977).). A Bonferroni post hoc analysis revealed that the APP_TS mice took significantly less time to locate the hidden platform during training days in comparison with APP (P≤ .0001), APP-GS (P≤ .0001), and APP-GS-TS (P≤ .0001) and APP-GS took significantly more time than the APP (P≤ .0001), APP-TS (P≤ .0001) and APP-GS-TS (P≤ .0001). The APP-TS mice swam the least distance compared to the APP-GS (P≤ .0001) and APP-GS-TS (P = .012) and APP-GS mice swam the longest distance compared to APP (P≤ .0001) APP-TS (P≡ .0001) and APP-GS-TS (P≤ .0001). APP-TS mice spent the highest amount of time in the target quadrant during the probe test compared to APP (P =.015) and APP-GS (P≤ .0001) and APP-GS mice spent the least amount of time in the target quadrant compared to APP-TS (P≤ .0001) and APP-GS-TS (P = .005).

### Balance Beam (BB) Test

The APP-GS mice were significantly slower to traverse the beam and made more slips than all of the other groups at both 2 and 6 months. TS significantly improved performance on both measures for both APP and APP-GS group (see Figure 3). The overall significant differences among all the four groups in latency at 2 months is F (3,57) = 8.289, P≤ .0001, □2 = .319, power = .989; at 6 months is F (3,50) = 21.543, P≤ .0001, □2 = .584, power = 1.000 and in number of foot slips at 2 months is F (3,57) = 2.955, P = .041, □2 = .143, power = .668; at 6 months is F (3,50) = 31.003, P≤ .0001, □2 = .669, power = 1.000.

**Figure 3.**
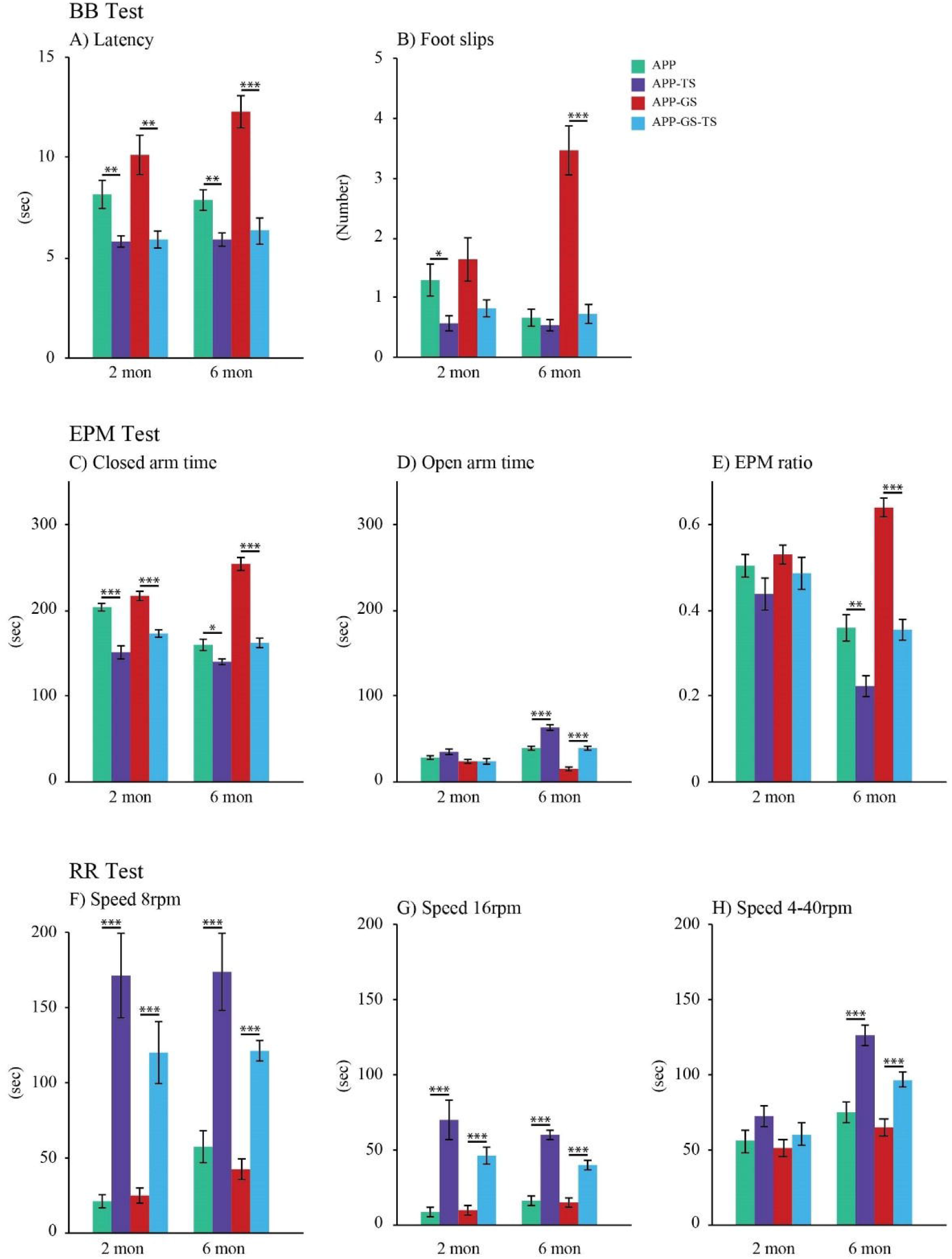
(A) The APP-TS mice were the fastest and APP-GS mice were the slowest in latency in both 2 and 6 months cohorts. (B) The APP-TS mice exhibited a decreased number of foot slips, whereas APP-GS exhibited an increased number of foot slips in both 2 and 6 months cohorts. (C) Shows significantly shorter closed arms time in APP-TS mice in both 2 and 6 months cohorts. (D) Shows significantly longer open arms time in APP-TS mice only in 6 months cohorts. (E) Shows highest EPM ratio for closed arms entries in APP-GS only in 6 months cohort. (F) An increased time spent with at (8 rpm) speeds in APP-TS mice of both 2 and 6 months cohorts. (G) An increased time spent at high (16 rpm) speeds in APP-TS mice of both 2 and 6 months cohorts. (H) There was no significant difference in the time spent at alternating (4-40 rpm) speeds among the groups in 2 months cohort, whereas APP-TS spent highest time in 6 months cohort.

At 2 months, APP-TS mice took significantly less time to cross the beam (F (1, 26) = 9.35, P = .005, □2 = .280, power = .835) and exhibited a reduced number of foot slips (F (1, 26) = 5.240, P = .031, □2 = .173, power = .594) compared to APP group. Similarly, the APP-GS-TS group took significantly less time to cross the beam (F (1, 31) = 11.427, P = .002, □2 = .283, power = .904), and exhibited a reduced but not significant reduced number of foot slips (F (1, 31) = 3.296, P = .080, □2 = .102, power = .419) relative to APP-GS mice at 2 months. A Bonferroni post-hoc analysis revealed that the APP_GS group took the longest time to cross the beam relative to APP-GS-TS (P = .001), and APP-TS (P≤ .0001), and had the highest number of foot slips relative to APP-GS (P = .046) at 2 months.

At 6 months, APP-TS mice took significantly less time to cross the beam (F (1, 25) = 9.684, P = .005, □2 = .296, power = .846) compared to APP group, but there was no difference in the number of foot slips (F (1, 25) = 0.639, P = .432, □2 = .027, power = .119). Similarly, the APP-GS-TS group took significantly less time to cross the beam (F (1, 25) = 26.156, P≤ .0001, □ 2 = .532, power = .998), and exhibited a significantly reduced number of foot slips (F (1, 25) = 28.868, P≤ .0001, □2 = .557, power = .999) relative to APP-GS mice at 6 months. At 6 months the APP-GS group took the longest time to traverse the beam relative to APP (P≤ .0001), APP-TS (P≤ .0001), and APP-GS-TS (P≤ .0001), and had a significantly a lower number of foot slips compared to APP (P≤ .0001), APP-TS (P≤ .0001), and APP-GS-TS (P≤ .0001) mice as per Bonferroni post hoc analysis.

### Rotarod (RR) test

The APP-TS group showed significantly better performance among all of the groups in all 8 rpm, 16 rpm, and 4-40 rpm at both 2 and 6 months age. TS significantly improved the RR performances in both APP and APP-GS mice at both at 2 and 6 months (see Figure 3).

At 2 months, the overall statistically significant effects in all four groups are at 8 rpm: F (3, 57) = 20.214, P≤ .0001, □2 = .534, power = 1.000; at 16 rpm: F (3, 57) = 16.991, P≤ .0001, □2 = .490, power = 1.000; and at 4-40 rpm: F (3, 57) = 1.688, P = .181, □2 = .087, power = .417. Similarly, at 6 months the overall statistically significant effects among the groups are at 8 rpm: F (3, 50) = 17.069, P≤ .0001, □2 = .527, power = 1.000; at 16 rpm: F (3, 50) = 41.693, P≤ .0001, □2 = .731, power = 1.000; and at 4-40 rpm: F (3, 50) = 18.794, P≤ .0001, □2 = .551, power = 1.000.

In the 2 months cohort, compared to APP mice, APP-TS group showed significantly improved performance in RR speeds, i.e., 8 rpm: F (1, 26) = 25.407, P≤ .0001, □2 = .514, power = .998; 16 rpm: F (1, 26) = 19.238, P≤ .0001, □2 = .445, power = .998. Similarly, a pairwise comparison revealed that APP-GS-TS mice exhibited improved performances in all three RR speeds, i.e., 8 rpm: F (1, 31) = 31.917, P≤ .0001, □2 = .524, power = 1.000; 16 rpm: F (1, 31) = 34.448, P≤ .0001, □2 = .543, power = 1.000 relative to APP-GS group. A Bonferroni post-hoc analysis revealed that the APP-TS group showed significantly improved performances among all the groups in two RR speeds, i.e., 8 rpm: APP (P≤ .0001) and APP-GS (P≤ .0001); 16 rpm: APP (P≤ .0001), APP-GS (P≤ .0001) at 2 months.

At 6 months, a pairwise comparisons revealed that APP-TS mice showed improved performances in all three RR speeds, i.e., 8 rpm: F (1, 25) = 18.343, P≤ .0001, □2 = .444, power = .984; 16 rpm: F (1, 25) = 92.328, P≤ .0001, □2 = .801, power = 1.000; and 4-40 rpm: F (1, 25) = 36.372, P≤ .0001, □2 = .534, power = .998 compared to APP. Similarly, APP-GS_TS mice exhibited improved performances in all three RR speeds, i.e., 8 rpm: F (1, 25) = 60.218, P≤ .0001, □2 = .724, power = 1.000; 16 rpm: F (1, 25) = 25.708, P≤ .0001, □2 = .528, power = .998; and 4-40 rpm: F (1, 25) = 18.377, P≤ .0001, □2 = .444, power = .984 relative to APP-GS at 6 months. At 6 months the APP-TS mice exhibited significantly improved performances in all three RR speeds, i.e., 8 rpm: APP (P≤ .0001) and APP-GS (P≤ .0001); 16 rpm: APP (P≤ .0001), APP-GS (P≤ .0001); 4-40 rpm: APP (P≤ .0001) and APP-GS (P≤ .0001) as per Bonferroni post-hoc analysis.

### Elevated Plus Maze (EPM) test

The APP-TS mice were significantly less anxious and spent significantly more time in the open arms, and less time in the closed arms compared to the other experimental groups. TS reduced the anxiety like behavior in both APP and APP-GS mice (see Figure 3). At 2 months, the overall significant effects among all of the four groups were: open arm time (F (3, 57) = 3.146, P = .033, □2 = .151, power = .699), closed arm time (F (3, 57) = 26.577, P≤ .0001, □2 = .601, power = 1.000), and EPM ratio (F (3, 57) = 1.731, P = .172, □2 = .089, power = .427). At 6 months, the overall effects among all of the groups were: open arm time (F (3, 50) = 54.430, P≤ .0001, □2 = .601, power = 1.000), closed arm time (F (3, 50) = 60.348, P≤ .0001, □2 = .797, power = 1.000), and EPM ratio (F (3, 50) = 38.645, P≤ .0001, □2 = .720, power = 1.000).

A pairwise comparison of the 2 months cohort revealed that the closed arm time was significantly higher in APP mice compared to APP-TS (F (1, 26) ≤ 37.225, P .0001, □2 = .608, power = 1.000), and in APP-GS mice relative to APP-GS-TS (F (1, 26) = 29.931, P≤ .0001, □2 = .508, power = 1.000). No significant effects were observed among the groups in either open arm time or EPM ratio at 2 months. Similarly, in the 6 months cohort the open arm time was significantly higher in APP-TS mice compared to APP (F (1, 25) = 39.539, P≤ .0001, □2 = .632, power = 1.000) and in APP-GS-TS mice relative to APP-GS (F (1, 25) = 38.167, P≤ .0001, □2 = .624, power = 1.000). The closed arm time was significantly lower in APP-TS mice relative to APP (F (1, 25) = 7.675, P = .011, □2 = .250, power = .756) and in APP-GS-TS mice compared to APP-GS (F (1, 25) = 64.640, P≤ .0001, □2 = .712, power = 1.000) at 6 months. In addition, the EMP ratio was significantly lower in APP-TS mice compared to APP (F (1, 25) = 9.791, P = .005, □2 = .299, power = .850) and in APP-GS-TS relative to APP-GS (F (1, 25) = 54.353, P≤ .0001, □2 = .715, power = 1.000) group at 6 months.

A Bonferroni post-hoc analysis revealed that APP-TS mice spent significantly more time in the open arms compared to APP-GS (P = .021) and significantly less time in the closed arms compared to APP (P≤ .0001) and APP-GS (P≤ .0001) in the 2 months cohort. However, in the 6 months cohort APP-TS mice spent significantly more time in the open arms relative to APP (P≤ .0001), APP-GS (P≤ .0001), and APP-GS-TS (P≤ .0001). In contrast, at 6 months APP-GS spent the most amount of time in the closed arms relative to APP (P≤ .0001), APP-TS (P≤ .0001), and APP-GS-TS (P≤ .0001) as per Bonferroni post-hoc analysis. In addition, at 6 months the EPM ratio was significantly lower in the APP-TS mice compared to APP (P = .007), APP-GS (P≤ .0001), and APP-GS-TS (P = .019) groups, but was significantly higher in the APP-GS mice relative to APP (P≤ .0001), APP-TS (P≤ .0001), and APP-GS-TS (P≤ .0001) group as per Bonferroni post-hoc analysis.

### Impact of TS on the amyloid-β (Aβ) plaque pathology

TS positively influenced the formation of Aβ by reducing the area of plaques (%) in both APP and APP-GS mice in both 2 months and 6 months groups (see Figure 4). Although there was a trend of increased Aβ plaque (%), there were no significant differences were observed between males and females. As a result, the data from both males and females were collapsed together. The area of Aβ plaques (%) was significantly smaller in all 6 brain sections and similar pattern of significant results were observed after combining all the sections together in both 2 months and 6 months groups. In the 2 months cohort, APP-TS mice showed significantly reduced total Aβ plaques (%) area compared to APP mice (F (1, 27) = 460.47, P = .0001, □2 = .947, power = 1.000). In addition, the total of Aβ plaques (%) area reduced in APP-GS-TS compared to APP-GS mice (F (1, 21) = 264.06, P = .0001, □2 = .930, power = 1.000). A Bonferroni post-hoc analysis revealed that APP-GS has the highest total Aβ plaques (%) area compared to APP (P = .0001), APP-TS (P = .0001), and APP-GS-TS (P = .0001).

**Figure 4.**
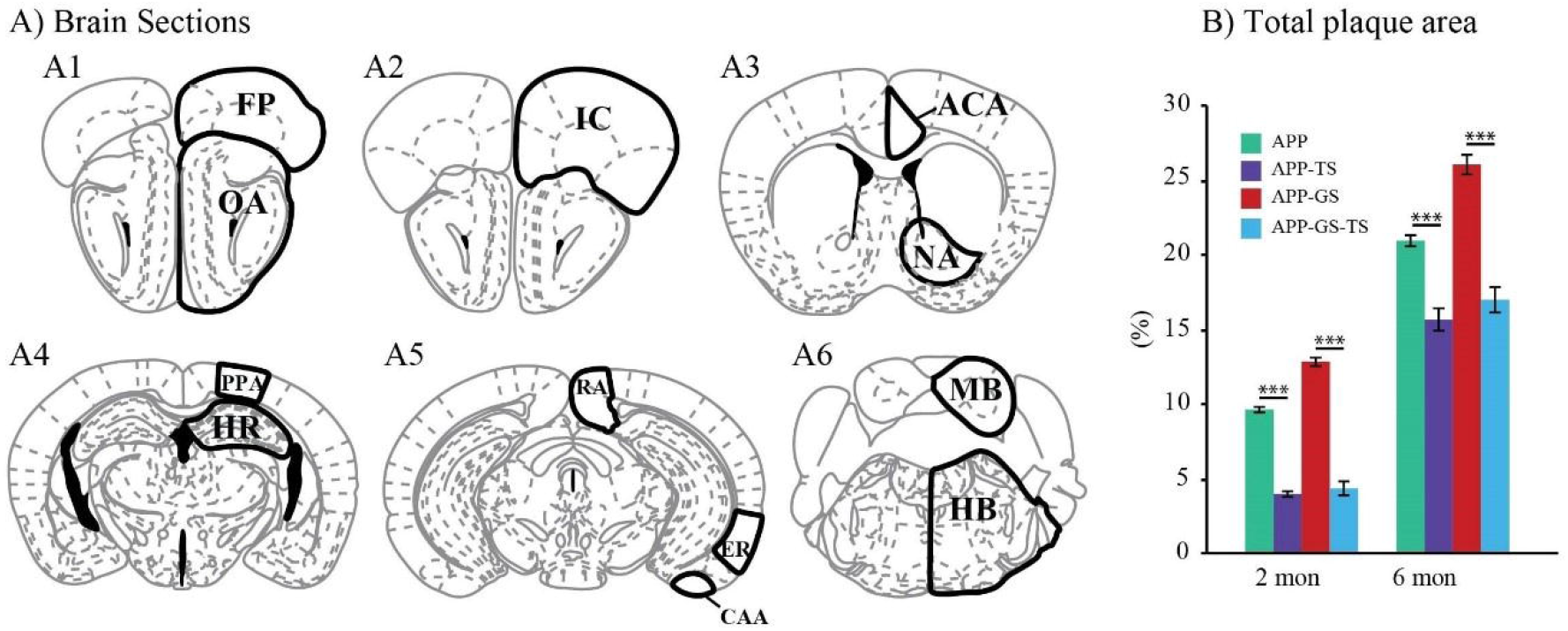
(A) Six coronal brain sections from A1-A6 with bregma position 3.2, 2.96, 0.98, −2.06, −3.08, and −5.34 mm. (B) Total Aβ plaque area (%): The APP-TS and APP-GS-TS groups had significantly lower plaque area (%) compared to APP and APP-GS mice, respectively.

In the 6 months cohort, APP-TS mice exhibited significantly reduced total Aβ plaques (%) area (F (1, 26) = 39.18, P = .0001, □2 = .610, power = 1.000) and APP-GS-TS mice showed significantly reduced total Aβ plaques (%) area compared to APP-GS mice (F (1, 26) = 71.88, P = .0001, □2 = .742, power = 1.000). Surprisingly, the total Aβ plaques (%) area is identical in both APP-TS and APP-GS-TS mice (P = 1.000) in both 2 months and 6 months cohorts. A Bonferroni post-hoc analysis revealed that APP-GS has the highest total Aβ plaques (%) area compared to APP (P = .0001), APP-TS (P = .0001), and APP-GS-TS (P = .0001).

TS reduced the formation of Aβ plaque area (%) in all the ROI’s in both 2 months and 6 months cohort (see Figure 5). In 2 months cohort, APP-TS mice showed significantly reduced Aβ plaque area (%) in IC (F (1, 27) = 11.860, P = .002, □2 = .313, power = .912), OA (F (1, 27) = 32.411, P =.0001, □2 = .555, power = 1.000), mPFC (F (1, 27) = 8.881, P = .006, □2 = .255, power = .818), ACA (F (1, 27) = 157.929, P = .0001, □2 = .859, power = 1.000), NA (F (1, 27) = 15.908, P = .0001, □2 = .380, power = .970), PPA (F (1, 27) = 19.427, P = .0001, □2 = .428, power = .989), HR (F (1, 27) = 21.520, P = .0001, □2 = .453, power = .994), EA (F (1, 27) = 83.513, P = .0001, □2 = .763, power = 1.000), CAA (F (1, 27) = 55.298, P = .0001, □2 = .680, power = 1.000), RA (F (1, 27) = 107.619, P = .0001, □2 = .805, power = 1.000), MB (F (1, 27) = 115.534, P = .0001, □2 = .816, power = 1.000), and HB (F (1, 27) = 1012.770, P = .0001, □2 = .975, power = 1.000) compared to APP mice. In addition, APP-GS-TS mice showed significantly reduced Aβ plaque (%) area in IC (F (1, 21) = 166.034, P = .0001, □2 = .892, power = 1.000), OA (F (1, 21) = 76.907, P = .0001, □2 = .794, power = 1.000), mPFC (F (1, 21) = 65.739, P = .0001, □2 = .767, power = 1.000), ACA (F (1, 21) = 101.076, P = .0001, □2 = .835, power = 1.000), PPA (F (1, 21) = 10.632, P = .004, □2 = .347, power = .873), EA (F (1, 21) = 62.500, P = .0001, □2 = .758, power = 1.000), CAA (F (1, 21) = 13.824, P = .001, □2 = .409, power = .942), RA (F (1, 21) = 103.406, P = .0001, □2 = .838, power = 1.000), MB (F (1, 21) = 828.071, P = .0001, □2 = .976, power = 1.000), HB (F (1, 21) = 2203.341, P = .0001, □2 = .991, power = 1.000) compared to APP-GS mice. Interestingly, no significant differences were observed in HR and NA between APP-GS and APP-GS-TS mice.

**Figure 5:**
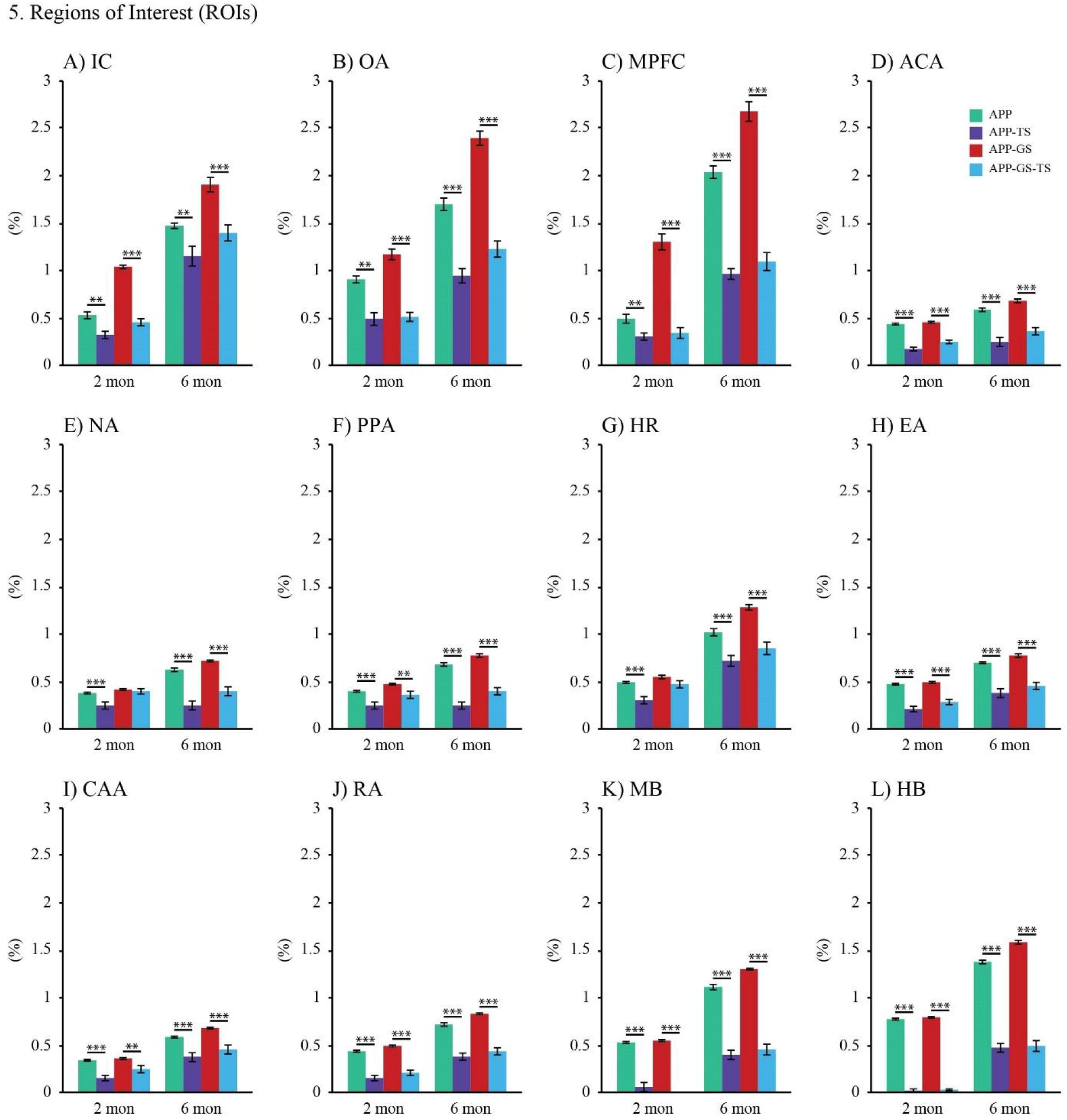
The Aβ plaque quantification in APP mice at the age of 2 and 6 months. The Plaque area (%) in 12 ROIs. (A-L) The APP-TS and APP-GS-TS has significantly lower plaque area (%) in all ROIs in both 2 and 6 months cohorts but the HR in 2 months cohort. HR, hippocampal region; IC, isocortex; mPFC, medial prefrontal cortex; NA, nucleus accumbens; OA, olfactory area. Results reported as mean ± S.E.M. Asterisks indicate P*≤ .05, p**≤ .01, and p***≤ .001.

Similarly in the 6 months cohort, APP-TS mice showed significantly fewer Aβ plaque (%) area in IC (F (1, 27) = 10.994, P = .003, □2 = .305, power = .890), OA (F (1, 27) = 49.394, P = .0001, □2 = .664, power = 1.000), MPFC (F (1, 27) = 144.628, P = .0001, □2 = .853, power = 1.000), ACA (F (1, 27) = 58.757, P = .0001, □2 = .702, power = 1.000), NA (F (1, 27) = 60.962, P = .0001, □2 = .709, power = 1.000), PPA (F (1, 27) = 153.605, P = .0001, □2 = .860, power = 1.000), HR (F (1, 27) = 23.404, P = .0001, □2 = .484, power = .996), EA (F (1, 27) = 58.530, P = .0001, □2 = .701, power = 1.000), CAA (F (1, 27) = 20.179, P = .0001, □2 = .447, power = .991), RA (F (1, 27) = 77.546, P = .0001, □2 = .756, power = 1.000), MB (F (1, 27) = 179.092, P = .0001, □2 = .878, power = 1.000), HB (F (1, 27) = 290.317, P = .0001, □2 = .921, power = 1.000) compared to APP mice. In addition, the APP-GS-TS showed significantly reduced Aβ plaques (%) area in IC (1, 25) = 20.254, P = .0001, □2 = .448, power = .991), OA (1, 25) = 110.124, P = .0001, □2 = .815, power = 1.000), mPFC (1, 25) = 125.393, P = .0001, ? 2 = .839, power = 1.000), ACA (1, 25) = 55.361, P = .0001, □2 = .689, power = 1.000), NA (1, 25) = 62.410, P = .0001, □2 = .714, power = 1.000), PPA (1, 25) = 88.599, P = .0001, □2 = .780, power = 1.000), HR (1, 25) = 39.235, P = .0001, □2 = .611, power = 1.000), EA (1, 25) = 66.656, P = .0001, □2 = .727, power = 1.000), CAA (1, 25) = 23.155, P = .0001, □2 = .481, power = .996), RA (1, 25) = 91.790, P = .0001, □2 = .786, power = 1.000), MB (1, 25) = 207.412, P = .0001, □2 = .892, power = 1.000), and HB (1, 25) = 358.998, P = .0001, □2 = .935, power = 1.000) in comparison to APP-GS mice in 6 months cohort.

## Discussion

The key findings of this study were: 1) early TS reduces the adverse effect of GS; 2) early TS reduces the AD-like behavioral symptoms; and, 3) early TS reduces the Aβ pathology in adult offspring at 2 and 6 months. We consider each finding in turn.

### Early TS mitigates the adverse effect of GS in the adult offspring at both 2 and 6 months

GS has a detrimental influence on brain and behavioural development. As per the previous publications on the effect of GS on adult offspring in our lab, it has been demonstrated that GS accelerates the progression of Aβ plaque deposition in adult APP offspring mice and the impaired cognitive, motor, and anxiety-like behaviours are associated with accumulation of Aβ plaque in the brain (Jafari et al., 2017; 2018; 2019). The behavioural findings from this study also demonstrate the similar deficits in cognitive, motor, and anxiety-like behaviours as a result of GS in the adult offspring. In this project we applied early TS as a therapeutic intervention with a goal of reversing the effect of GS in adult APP offspring. Our results demonstrate that it was successful.

GS influences the HPA and hypothalamic-pituitary-gonadal (HPG) axes and these two axes also modify the function of each-other (Toufexis et al. 2014). GS elevates the corticosterone levels (Jafari, et al., 2017) in dams, which initiates a long-lasting dysfunction in the HPA negative feedback loop. Elevated corticosterone in the dams influences the level of corticosterone in the offspring and adversely impacts their brain and social stability (Saavedra-Rodriguez & Feig, 2013). Stress during pregnancy also affects the prefrontal cortex and hippocampus development of the offspring and the alterations of these are associated with reduced spatial and cognitive functions (Mychasiuk et al., 2011). As a result, the offspring also exhibit impaired cognition and motor skills and anxiety-like behaviors as adults. In our study, the APP mice that received GS spent less time in the open arms and had fewer entries to the open arms of the maze compared to the APP mice. In addition, APP-GS mice showed less exploratory behaviour towards the novel object. Both APP and APP-GS groups that received TS exhibited improved performances in both EMP and NOR tests. Mice that received TS showed less anxiety and more interest in novel object exploration. GS also impaired cognitive and motor skills in adult offspring. The results also demonstrated that TS improved the performances of adult offspring in the MWT, BB, and RR test. Research on rats has demonstrated that TS early in life reduced the corticosterone level in infants (Jutapakdeegu et al., 2003). Studies on rats also showed that TS increased FGF-2 in skin and brain (Gibb, 2004; Gibb et al., 2021). TS also increases BDNF, which enhances the proliferation and protection of neurons in the hippocampus (Antoniazzi et al, 2016), and perhaps other brain regions such as visual cortex (McAllister et al, 1995), thalamus, hypothalamus, striatum, and septum (Pencea et al., 2001) as well. Hence, TS during earlier life stages plays a critical role in neuroplasticity in later in life.

Surprisingly, we observed increased old object time in the NOR test in both of the APP and APP-GS offspring that received TS. This may be a result of enhanced exploration and activity in mice that received TS. Another important observation was that the effect of TS was stronger in the 6 months cohort than in the 2 months cohort likely because the earliest onset of AD-like symptoms and pathology is at about 3 months in this APP mouse model (Jafari et al., 2017).

The symptoms of AD in the APP mice increases as the disease progresses with increased age. The application of TS during the earlier stages of their life plays an important role by prolonging the onset of AD and also improving exploratory, spatial, and anxiety-like behaviours in both 2 and 6 months adult offspring.

### Early TS improves the AD-like symptoms in adult offspring at 2 and 6 months

Impaired learning and memory along with deterioration in motor skills and elevated anxiety are common symptoms of AD in humans. A series of studies in our lab have shown the similar deficits in APP mice (Jafari, et al., 2017, 2018, 2019; Mehla et al., 2019; Karem et al., 2021). In addition, the declined cognitive and motor skills are associated with the formation of Aβ plaques and with age as the formation of Aβ plaques increases with age in these APP mice (Jafari et al., 2018; Mehla, et al., 2019). The goal of this study was to see whether application of early TS would influence learning and memory, motor, and anxiety-like behaviours in adult APP offspring at 2 and 6 months. The results from the NOR and MWT tests suggest that TS improves the cognition in the mice that received TS. In the NOR test, APP mice that received TS demonstrated higher preference for the novel object. Similarly, in the MWT test the APP mice that received TS displayed significantly shorter latency and swim distance during the training days, and significantly longer probe time, suggesting improved spatial learning and memory (see also Angeles et al., 2016; Kolb & Gibb, 2010).

A positive impact of TS also was observed in motor skills among APP adult offspring at both 2 and 6 months. In the BB test, the APP mice that received TS were faster to traverse the beam and had fewer slips. Similarly, in the RR test APP mice that received TS were able to stay longer time in the rotating rod in all three different speeds at both 2 and 6 months. The findings from both BB and RR tests suggest that improvements in motor balance and coordination in APP mice result from early TS. Research with APP mice has also shown that age plays a vital factor in deteriorating AD-like symptoms in that the motor deficits in APP mice and at 6 months are worse than at 2 months (Jafari et al., 2018). In this study, we showed that early TS prevents the deficits in motor skills even at 6 months of age in APP mice.

In addition, TS significantly reduced anxiety-like behaviour in both control and APP mice. In the EPM test, the mice that received TS spent the highest time in the open arms and exhibited the highest EPM ratio for the open arms. These results suggest that early TS helped to reduce the anxiety-like behaviour in the APP mice, and these findings were greater at 6 months. The mechanisms of the effects of TS on both AD and stress are not firmly established and likely are multiple including epigenetic changes as well as changes in neuronal morphology.

### Early TS reduces the Aβ pathology in adult offspring at 2 and 6 months mice

Aβ plaques are one of the vital hallmarks of the neural symptoms of AD. In this study, we have shown that early TS reduces the development of Aβ plaques in APP mice, as well as APP-GS mice in 2 months and 6 months cohorts. The positive influence of early TS has been observed in almost all ROIs we have analyzed. Although the formation of Aβ plaques is visible throughout neocortex and hippocampal area, it is well established that Aβ plaques begin to develop in OA (Wesson et al., 2010), EA (Sipos et al., 2007), and RA (Poirier et al., 2014) brain regions first, then progressively spread to rest of the brain regions. Our findings demonstrate that TS significantly reduced the Aβ pathology in all ROIs such as IC, OA, mPFC, NA, PPA, HR, CAA, EA, RA, MD, and HB. The correlation of stress and AD-like dementia and Aβ pathology is well established in human (Justice, 2018) as well as in APP mice (Zahra et al., 2017; 2018; 2019). Our investigation also shows that early TS significantly reduces Aβ pathology in APP and APP-GS mice and the reduction of Aβ plaques is correlated with cognitive, motor, and anxiety-like behaviour. Now, the question is how exactly TS positively influences the development of Aβ pathology and improves the AD-like symptoms. There is no known neural mechanism of TS yet although it may be related to the increase in FGF-2 or other epigenetic effects.

The influence of TS on the endocrine system is well known. TS is applied to the largest human organ, skin, which contains growth factors and receptors for peptide hormones and neurotransmitters. Out of 22 growth factors, FGF-2 plays a vital role in neuronal growth. Reduction of Aβ plaques also increases the changes of neuronal communication in inter- and intra-brain regions which supposedly helps with AD like symptoms (Spires-Jones and Hyman, 2015). One very important observation in our study is that early TS reduces the Aβ formation in APP-GS mice almost similar to APP-TS mice in both 2 months and 6 months cohort separately. Therefore, we can conclude that the early TS has a positive influence APP-GS mice as well as APP mice. There is evidence that shows that stress alters the HPA (Hypothalamus-Pituitary-Adrenal)-axis by increasing corticosterone, epinephrine, and norepinephrine (Costa et al., 2020) and these authors also showed that TS positively reversed the impact of stress. Although no clear mechanisms of action of TS has been established yet, studies show application of TS during neonatal period (Freitas et al., 2015) or adult (Roversi et al., 2019) reduces corticosterone and adrenocorticotrophic hormone in adult rats. Research also shows that early TS also enhances the spine density and dendritic connectivity on medial pre-frontal cortex (mPFC) and orbital frontal cortex (OFC) and amygdala in healthy adult rats (Richards et al, 2012), as well as, in a valproic acid rat model of autism (Raza et al., 2015). Neonatal tactile stimulation also proven to positively impact the genetic generalized epilepsy by reducing the number and length of spike-wave discharges, a hallmark of absence epilepsy in adult Wister Albino Glaxo from Rijswijk (WAG/Rij) rats (Balikci et al., 2020).

The imbalance of excitatory and inhibitory functions in the brain is another theory of many neurological disorders along with AD. It is well established that Aβ plaques reduce neural network connectivity by decreasing synaptic glutamatergic transmission (Sinnen et al., 2016) and GABAergic interneurons connectivity in APP mice (Verret et al., 2012). Early TS increases the GABAA receptor binding levels in prefrontal cortex, hippocampus, and amygdala rats (Caldji et al., 2003). Another important aspect of AD pathology is that dopaminergic neuron degeneration is observed in ventral tegmental area but not in substantia nigra in the Tg2576 mouse model of AD; and the compromised dopamine projection to nucleus accumbens and hippocampus results in impaired synaptic plasticity in CA1 region (Nobili et al., 2017). No study was found that shows a positive impact of TS on hippocampal dopaminergic projections. However, Maruyama et al. (2012) showed that application of bilateral TS for 5 min increased dopamine release in nucleus accumbens in both conscious and anesthetised adult rats.

Brain-derived neurotrophic factor (BDNF), which is a member of the neurotrophin family of growth factors found in both peripheral and central nervous system, plays vital role in neuronal plasticity, neurotransmitter modulation, and neuronal survival and growth. Reduced BDNF mRNA and protein levels were observed in the frontal cortex and hippocampus in post-mortem brain samples from end-stage AD patients (Ferrer et al., 1999), in the parietal cortex from an early stage of AD patients (Peng et al., 2005), and in the temporal cortex of AD patients (Lee et al., 2005). An *in vitro* study shows that Aβ plays a role in reducing BDNF level (Zheng et al., 2010), which may lead to impaired synaptic plasticity. Intravenous infusion of BDNF with ADTC5, a blood brain barrier modulator, increased the level of BDNF in the hippocampus of APP/PS1 mice and increased BDNF was associated with higher efficacy in NOR and Y-maze tasks (Kopec et al., 2020). Neonatal TS from day 8-14 has proven to be beneficial in increasing BDNF level, and TS from day 1-7 and day 15-21 hasve been shown to reduce glucocorticoid receptors in the hippocampus in adult rats (Antoniazzi et al., 2016). An EEG study showed that visuo-tactile stimulation increases serum BDNF expression level in humans and this increased BDNF level is correlated with gamma oscillations in the left parietal cortex (Hiramoto et al., 2017). FGF-2 plays very important role in neuronal proliferation, differentiation, and neurogenesis (Woodbury and Ikezu, 2014). Reduced FGF-2 was found in post-mortem brains in AD (Takami et al., 1998). Subcutaneously administered FGF-2 in APP23 mice has been shown to reduce Aβ plaques and tau tangles by reducing the expression of BACE1, an enzyme that plays a role for Aβ production (Katsouri et al., 2015). TS on healthy adult rats showed increased FGF-2 in the striatum and hippocampus, but did not show any benefit to hemiparkinsonian adult rats (Effenberg et al., 2014).

## Conclusion

In the current study we were able to show that early TS helped to reduce the adverse effect of gestational auditory stress in APP adult offspring as well. A concurrent study showed that application of TS in adult APP mice reduces AD-like behavioral symptoms and pathology in adult APP mice (Hossain et al, 2021). These results suggest that TS has a preventative mechanism in this APP mouse model of AD. Further research is required to discover the neural mechanism regarding change in the gene expression, electrophysiology, neurotransmitters, FGF-2, and synapses as a result of early TS. In addition, we have only investigated the Aβ pathology in this study but we still need to explore whether early TS would influence cerebral inflammation, the tau-related tangles, and cerebral cholinergic activity in the AD mouse model.

## Acknowledgements

This work was supported by Natural Sciences and Engineering Research Council of Canada (NSERC) Discovery Grant #40352 to MHM, Alberta Innovates (MHM), Alberta Alzheimer Research Program (MHM), Alzheimer Society of Canada (MHM), Alberta Prion Research Institute (MHM), Canadian Institute for Health Research (MHM), and Alberta Registered Nurse Education Trust (SRH). We thank Dr. Takashi Saito and Prof. Takaomi C Saido from Laboratory for Proteolytic Neuroscience RIKEN Center for Brain Science, Wako-shi, Saitama, Japan” for providing the App^NL-G-F/NL-G_F^ mice as a gift. We also thank Di Shao for animal breeding.

Writers would like to acknowledge the grant supported by

## Authors Contribution

S.H., Z.J., M.H.M., and B.E.K. designed and conceptualized the study. M.H.M., and B.E.K. supervised the study. S.R.H. performed the behavioural experiments. S.R.H. analyzed the behavioural data. S.R.H., and H.K. performed the immunohistochemistry. H.K., analyzed the immunohistochemistry data. S.R.H., and B.E.K. wrote the manuscript. S.H., Z.J., M.H.M., and B.E.K. all commented on and edited the manuscript.

